# Comparison of odor responses of homologous medial and lateral glomeruli mapped in the olfactory bulb of the mouse

**DOI:** 10.1101/643288

**Authors:** Tokiharu Sato, Ryota Homma, Shin Nagayama

## Abstract

Olfactory sensory neurons expressing same-type odorant receptors typically project to a pair of glomeruli in the medial and lateral sides of the olfactory bulbs (OBs) in rodents. However, their functional properties remain unclear, because the majority of medial glomeruli are hidden in the septal OB. Recently, trace amine-associated odorant receptors were identified that project to a pair of glomeruli uniquely located in the dorsal OB. We measured the odorant-induced calcium responses of these glomeruli simultaneously and found that they exhibited similar temporal response patterns. However, the medial glomeruli had significantly larger respiration-locked calcium fluctuations than the lateral glomeruli. This trend was observed with/without odorant stimulation in postsynaptic neurons but not in presynaptic sensory axon terminals. This indicates that the medial rather than the lateral OB map enhances the respiration-locked rhythm and transfers this information to higher brain centers.

**Impact Statement:** This study used *in vivo* calcium imaging to document the odor-evoked responses in paired glomeruli, demonstrating that activation in medial glomeruli more strongly impacts respiratory-linked odor processing.

## Introduction

Parallel processing of multiple streams of information improves the speed of processing and provides redundancy for fail-safe operations. Biological parallel streams of information in the brain are not typically identical neuronal circuits but have unique as well as common properties (Kandel et al., 2013). How the brain organizes the distinct processing streams and combines them is not well understood.

Odor information is represented as spatial/temporal glomerular activity patterns on the surfaces of the olfactory bulbs (OBs). In rodents, olfactory sensory neurons (OSNs) expressing the same types of odorant receptors, among ~1,000 repertoires (Buck, 1996), convert chemical signals into electrical signals in a respiratory rhythm and project to approximately two glomeruli in the OB: one on the medioventral side and the other on the dorsolateral side. The axons of these OSN projections traverse the medial/septal and lateral surfaces of the OBs, respectively. Because the two homologous glomeruli are arranged symmetrically in the OB, odor information is represented and processed in two mirror maps (Mombaerts et al., 1996; Nagao et al., 2002, 2000; Zapiec and Mombaerts, 2015). The paired glomeruli are connected via axon collaterals of tufted cells in a point-to-point manner (Belluscio et al., 2002; Lodovichi et al., 2003; Marks et al., 2006). Although the anatomical arrangements of the two maps have been studied, less is known about the functional connections between them (Zhou and Belluscio, 2012, 2008). One pivotal idea is that the complex structure of the olfactory epithelium (OE) affects the sensitivity and timing of odorant responses of OSNs such that a delayed odorant response in one of the maps occurs with low odor concentrations (Kimbell et al., 1997; Schoenfeld and Cleland, 2006; Zhao et al., 2006; Zhou and Belluscio, 2012). The similarity in the spatiotemporal patterns representing odor information between these glomeruli remains unknown, partly because of the inaccessibility of the medial map, but is essential to understand the features that are common and uncommon between the two streams.

Trace amine-associated receptors (TAARs) were recently recognized as a second group gene family of odorant receptors (Liberles and Buck, 2006). Pairs of OSNs expressing TAARs project to glomeruli in the mediodorsal OB (Dewan et al., 2013; Liberles, 2015; Pacifico et al., 2012; Zhang et al., 2013), which can frequently be observed simultaneously. Moreover, these two glomeruli are functionally identifiable because of their highly selective responses to the specific odorant at a low concentration (Zhang et al., 2013). In the present study, we measured simultaneous odorant responses of these homologous glomerular pairs and compared the response properties between medial and lateral maps. These glomeruli exhibited similar activity patterns of response onset latency, rise and decay times, and amplitudes. However, the medial glomeruli showed significantly larger respiration-locked fluctuations than the lateral glomeruli. The difference was observed in postsynaptic neuronal responses but not presynaptic terminal activity, suggesting that despite similar inputs, the medial map neurons and/or circuits enhance the respiration-locked activity for further odor information processing.

## Results

### Expression patterns of GCaMP3 in OBs from Cre mouse driver lines

We recorded odor-evoked neuronal activity in the OBs of mice from multiple transgenic mouse lines. To express the genetically encoded calcium indicator GCaMP3 (Tian et al., 2009) in different types of neurons in the OB, we used four Cre recombinase (Cre)-driver mouse lines. Specifically, OMP-Cre (Li et al., 2004), Gad2-Cre (Taniguchi et al., 2011), DAT-Cre (Bäckman et al., 2006), and Pcdh21-Cre (Nagai et al., 2005) mouse lines were crossed with a Cre-inducible GCaMP3 reporter mouse line (Ai38) (Zariwala et al., 2012) so that GCaMP3 was expressed in OSNs and GABAergic, dopaminergic, and mitral/tufted cells, respectively. We first verified the GCaMP3 expression pattern in each of the Cre-driver mouse lines (Fig. 1 A–D). In OMP-Cre mice, GCaMP3 signals were clearly detected only in OSNs in the glomerular layer (GL) (Fig. 1A). Consistent with a previous report (Wachowiak et al., 2013), GCaMP3 in Gad2-Cre mice was strongly expressed in the external plexiform layer (EPL) and in the granule cell layer (GCL); the expression was predominantly by granule cells, but enhanced detection methods also revealed expression by periglomerular cells in the GCL (Fig. 1B). In DAT-Cre mice, GCaMP3 signals were mostly restricted to the GL (Fig. 1C). High magnification of this layer indicated that GCaMP3 was expressed by juxtaglomerular cells, which were considered to be short-axon (SA) cells (Kiyokage et al., 2010) (inset of Fig. 1C). This pattern of expression was similar to that in the TH (tyrosine hydroxylase)-Cre line, another driver line for expression in dopaminergic cells (Wachowiak et al., 2013). In Pcdh21-Cre mice, GCaMP3 was specifically expressed in mitral/tufted cells, as GCaMP3 signals appeared in somata and neurite processes in superficial EPL and the mitral cell layer (MCL) (Fig. 1D), as reported previously (Huang et al., 2013; Mizuguchi et al., 2012; Nagai et al., 2005). Although reporter expression is induced in OSNs in another Pcdh21-Cre-driver line (Wachowiak et al., 2013), we did not observe this ectopic expression in our mice. In summary, these Cre mouse lines exhibited the expected cell-type-specific GCaMP3 expression patterns.

**Figure 1:**
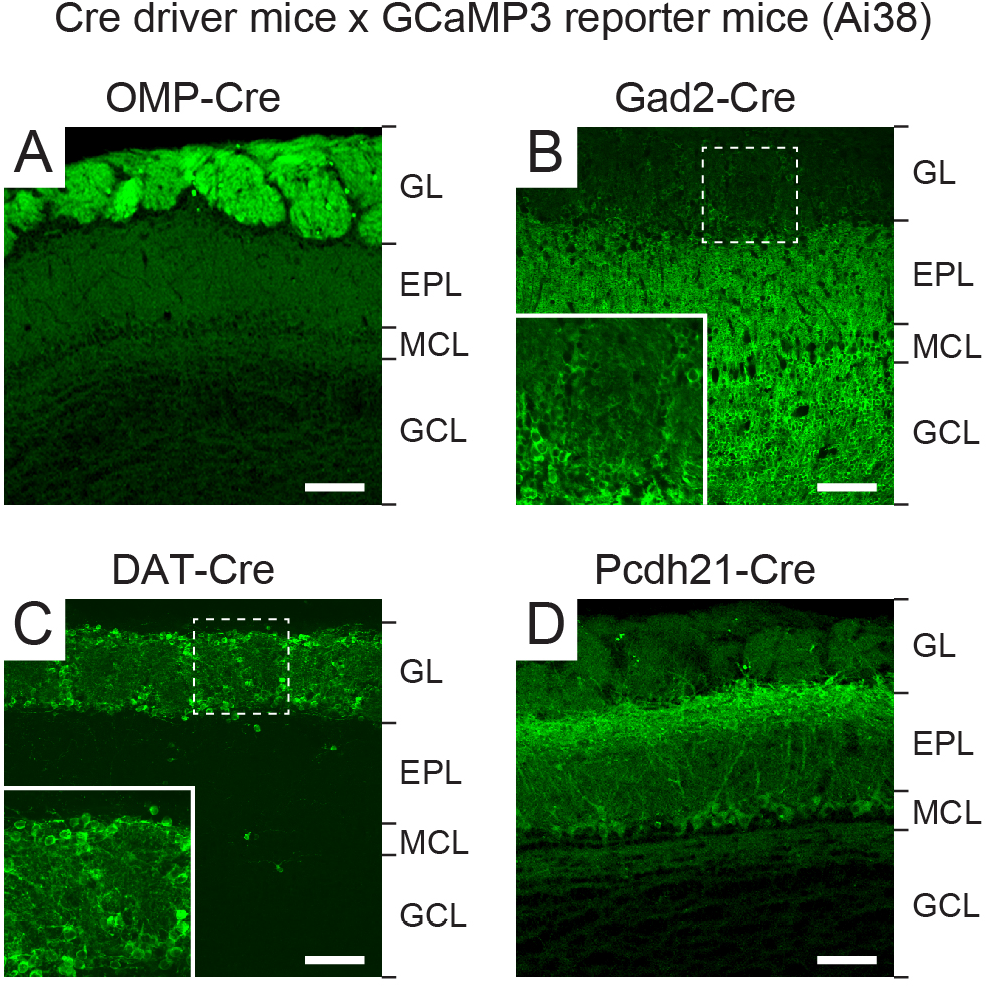
Cell-type-specific expression of GCaMP3 in OBs of mice from different transgenic lines. (A–D) Confocal images of OBs in Cre-dependent GCaMP3 reporter mice crossed with OMP-Cre (A), Gad2-Cre (B), DAT-Cre (C), and Pcdh21-Cre (D) mice. Magnified views of the dashed squares are shown in the insets in B and C. GL, EPL, MCL, and GCL indicate glomerular layer, external plexiform layer, mitral cell layer, and granule cell layer, respectively. Scale bars, 100 μm.

### Identification of homologous glomeruli in medial and lateral maps

*In vivo* optical imaging of the dorsal OB has not been utilized as a means to record activity in the medial map, because most of the glomeruli are located in the medioventral region of the OB (Inaki et al., 2002; Johnson et al., 2009, 2005, 2004, 1999, 1995, Johnson and Leon, 2000, 1996; Mori et al., 2006; Nagao et al., 2000, 2002; Taniguchi et al., 2003; Zapiec and Mombaerts, 2015). However, it was recently demonstrated that OSNs expressing TAARs project to two or a few glomeruli in caudal regions of the dorsal OB (Dewan et al., 2018, 2013; Johnson et al., 2012; Liberles, 2015; Pacifico et al., 2012; Zhang et al., 2013). These medial and lateral glomeruli can easily be identified, because axons from TAAR-expressing OSNs projecting to the dorsomedial glomeruli transverse the anteromedial (septal) surface of the OB, whereas those projecting to the dorsolateral glomeruli transverse the anterolateral surface. The axonal trajectories and odorant response profiles of glomeruli receiving projections from OSNs expressing TAAR3 and TAAR4 are well characterized (Dewan et al., 2018, 2013; Pacifico et al., 2012; Zhang et al., 2013). Specifically, these glomeruli are highly sensitive to isopentylamine (IPA) and phenylethylamine (PEA), respectively. Therefore, we imaged simultaneously these medial and lateral glomeruli that receive input from TAAR3- and TAAR4-expressing OSNs.

Using IPA and PEA at final concentrations of 0.02% and 0.002%, which reliably and specifically induce activation of the glomeruli receiving inputs from TAAR3- and TAAR4-expressing OSNs, respectively (Dewan et al., 2018, 2013; Pacifico et al., 2012; Zhang et al., 2013), we identified distinct pairs of glomeruli comprising OSNs and GABAergic, dopaminergic, and mitral/tufted cells in the caudal areas of the dorsal OB (Fig. 2A and Fig. 3A). The locations of these homologous glomeruli are consistent with the positions of glomeruli receiving inputs from TAAR3- and TAAR4-expressing neurons in previous reports (Dewan et al., 2018, 2013; Pacifico et al., 2012; Zhang et al., 2013).

**Figure 2:**
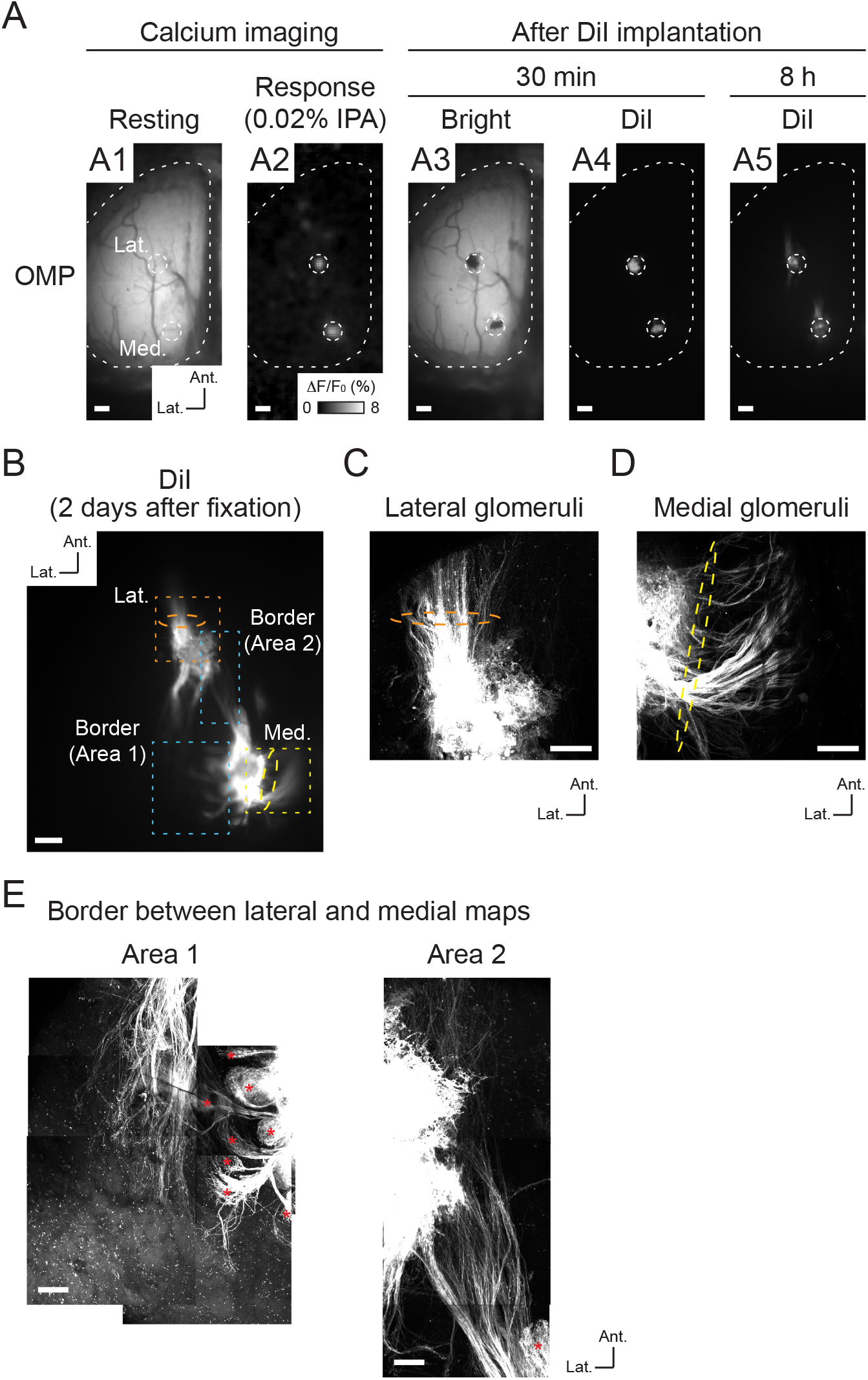
OSN axon trajectories for IPA-responsive glomeruli. (A) Process for the DiI labeling of OSN axons. A1, resting fluorescence of GCaMP3 in the dorsal OB of OMP-Cre mouse; A2, IPA (0.02%)-responsive homologous glomeruli were observed by calcium imaging (color scale indicates ΔF/F0 [%] of GCaMP3 signal); A3, brightfield image after DiI implantation; A4, DiI fluorescence 30 min after DiI implantation; A5, DiI fluorescence 8 h after DiI implantation. The locations of IPA-responsive glomeruli are indicated by the white dotted circles. (B) Two-photon microscopy image of DiI-labeled glomeruli and OSN axons 2 days after DiI implantation. (C) Magnified image of area denoted by orange dotted square in B associated with lateral glomerulus. (D) Magnified image of area denoted by yellow dotted square in B associated with medial glomerulus. Two-photon microscopy images of OSN axons that transverse the lateral and medial surface of OB; orange and yellow dotted ellipses (in B, C, and D) represent major axonal projections from lateral and medial glomeruli. (E) Two-photon microscopy images of areas of lateral/medial border denoted by blue dotted square in B. Red asterisks indicate axon termination in several glomeruli in which the labeled OSNs probably passed through the surface of the DiI-implanted glomeruli. Some minor axons which did not show clear axon terminations in a glomerulus were observed in area 2 in this case. Ant., anterior. Lat., lateral. Scale bars, 200 μm in A and B, 50 μm in C and D, 100 μm in E.

**Figure 3:**
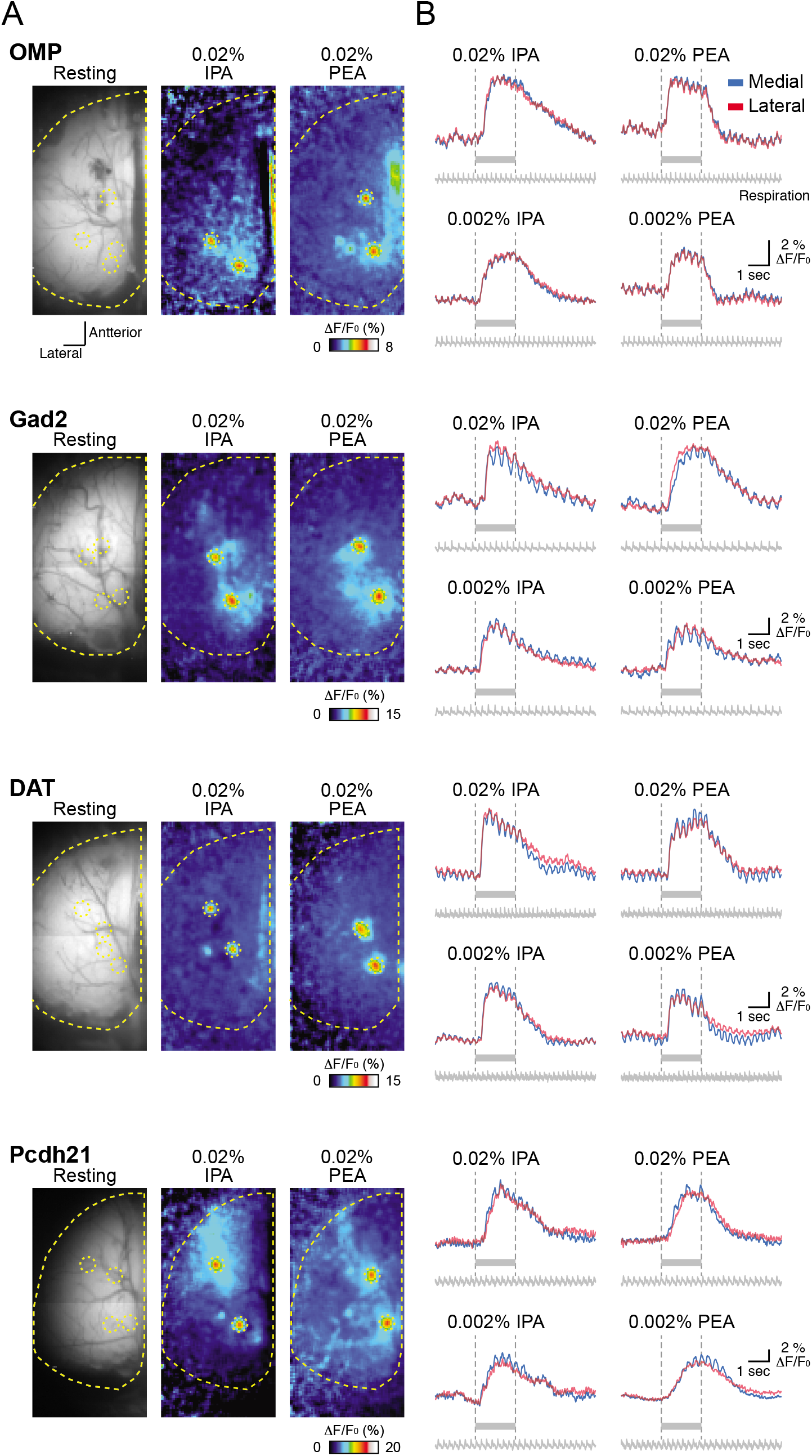
Odor-evoked response maps and traces among different types of OB neurons. Odor-evoked response maps (A) and traces (B) from OMP-Cre, Gad2-Cre, DAT-Cre, and Pcdh21-Cre mice. Resting GCaMP3 fluorescence images are displayed in the left panels in A. Pseudocolored images in the middle and right panels in A indicate responses to 0.02% IPA and PEA, respectively. Yellow dotted lines and circles in each image show approximately the edge of the left OB and homologous glomeruli evoked by 0.02% IPA and PEA, respectively. The color scales represent ΔF/F0 (%) of GCaMP3 signal. Traces shown in B represent GCaMP3 fluorescence changes from two subsets of homologous glomeruli evoked by indicated concentrations of IPA and PEA shown in A. Red and blue traces indicate lateral and medial glomeruli, respectively. Gray bars under each trace and vertical dotted lines indicate the timing of odor stimulation. Respiration signals are also shown under the traces.

To confirm that the glomeruli pairs represent medial and lateral maps, we retrogradely labeled TAAR3-expressing OSN axons by implanting the IPA-responsive glomeruli in OMP-GCaMP3 mice with DiI (1,1’-dioctadecyl-3,3,3’3’-tetramethylindocarbocyanine perchlorate) crystals. Optical observation performed 8 h later revealed labeling of OSN axons in both lateral and medial glomeruli (Fig. 2A5). These labeled axons were observed more clearly in fixed tissue 2 days later (Fig. 2B). Dorsolateral to the lateral IPA-responsive glomerulus, the majority of the labeled axons were oriented in an anterolateral direction (orange dashed ellipses in Fig. 2B and C). By contrast, the majority of axons dorsomedial to the medial IPA-responsive glomerulus were in an anteromedial direction (toward the septum) (yellow dotted ellipses in Fig. 2B and D). Notably, multiple glomerular structures were revealed by the terminal branches of labeled axons in the medial region (red asterisks in Fig. 2E), indicating that these axons originated from the medial side. Taken together, these results indicate that the pairs of recorded glomeruli were homologous pairs representing both lateral and medial maps. Therefore, our experimental design provides a unique opportunity to record simultaneously the odor-evoked neuronal activity of homologous glomeruli in the medial and lateral maps in OBs.

### Similar temporal odor representations between medial and lateral maps in the OB

The medial and lateral glomeruli pairs represent inputs from odorant receptors of the same type within medial and lateral regions, respectively, of the complex OE structure. Receptors in these regions may be exposed to different air flow rates and mucosal volumes. We hypothesized that these differences would be reflected in the timing of the glomeruli responses. However, the calcium signals from all cell types examined in medial and lateral glomeruli in response to IPA and PEA had similar amplitudes and temporal patterns (Fig. 3B). Further analyses revealed that the timing of odor inputs to both maps was similar, as revealed by the onset latency measured as the time at which the calcium signal exceeded the threshold from first inhalation during odor stimulation (Fig. 4). The rise times of the responses, which are an indicator of response speed and reflect the neuronal spike frequency, were similar between medial and lateral glomeruli (Fig. 5). The rise time was assessed as the duration for the calcium signal to increase from 20% to 80% of the peak signal. Conversely, the similar decay times, during which the calcium signal decreased from 100% to 50% of the peak signals, suggested that the activity in one glomerulus was not prolonged relative to the other after the odor stimulus was turned off (Fig. 6). Both glomeruli in the pairs had responses that were similar in strength, as indicated by the peak amplitudes (Fig. 7). Overall, there were no significant differences between paired glomeruli in onset latency, rise time, decay time, and peak amplitude of the calcium responses in any of the cell types studied (two-tailed paired *t* tests, see Table 1).

**Figure 4:**
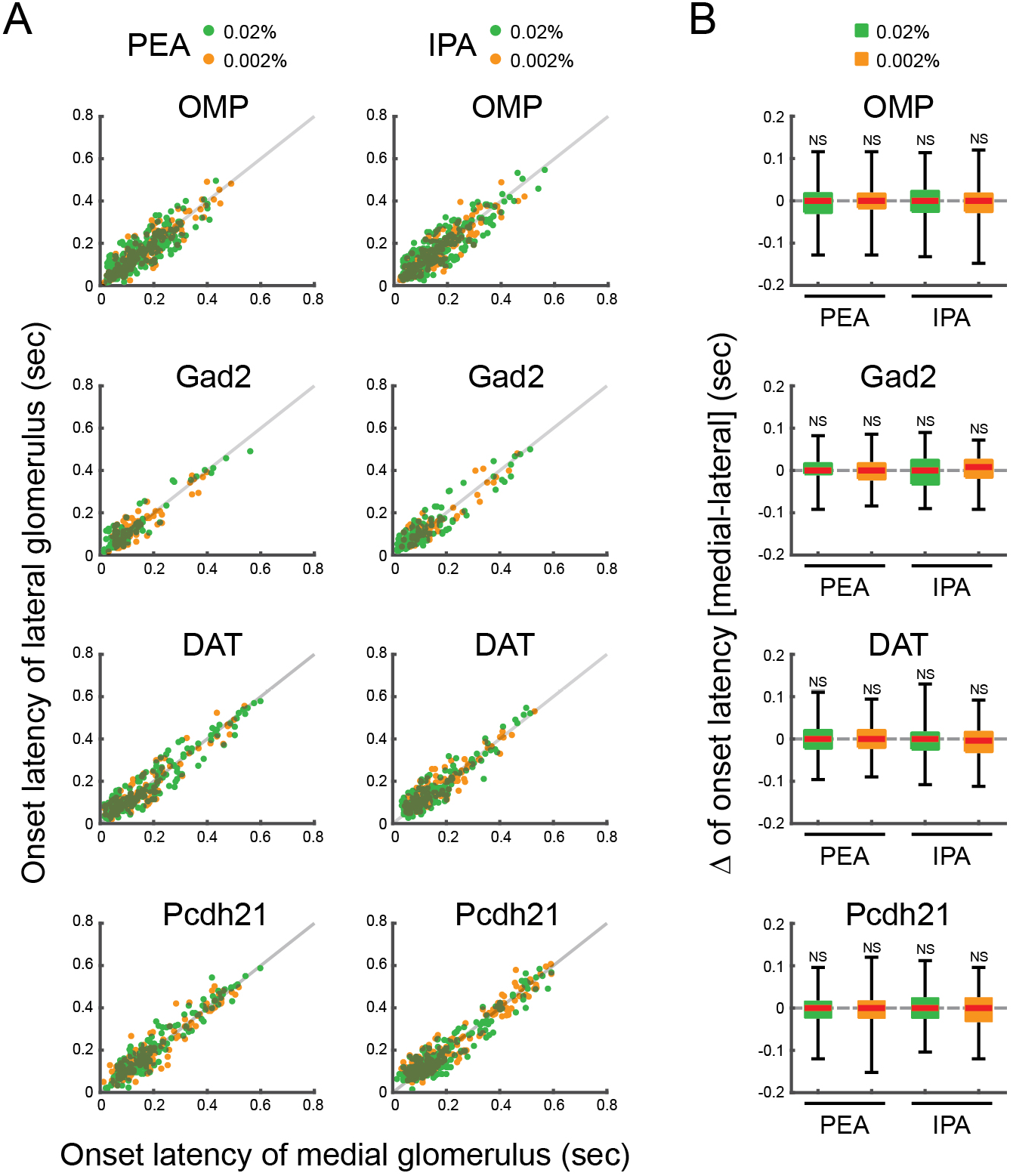
Onset latencies of medial and lateral glomeruli. (A) Scatter plots displaying the distributions of onset latencies of medial and lateral glomeruli, which responded to PEA and IPA stimuli. x and y axes indicate onset latencies of medial and lateral glomerular responses, respectively. Individual green and orange dots indicate single trial data of 0.02% and 0.002% PEA or IPA. Gray lines indicate the equal onset latency time points of medial and lateral glomerular responses. (B) Box plots displaying the distributions of the differences of onset latencies between medial and lateral glomeruli responses to 0.02% and 0.002% PEA or IPA. Red horizontal lines in each box indicate the medians. Quartiles are shown as whiskers. NS, not significant (two-tailed paired Student’s t test).

**Figure 5:**
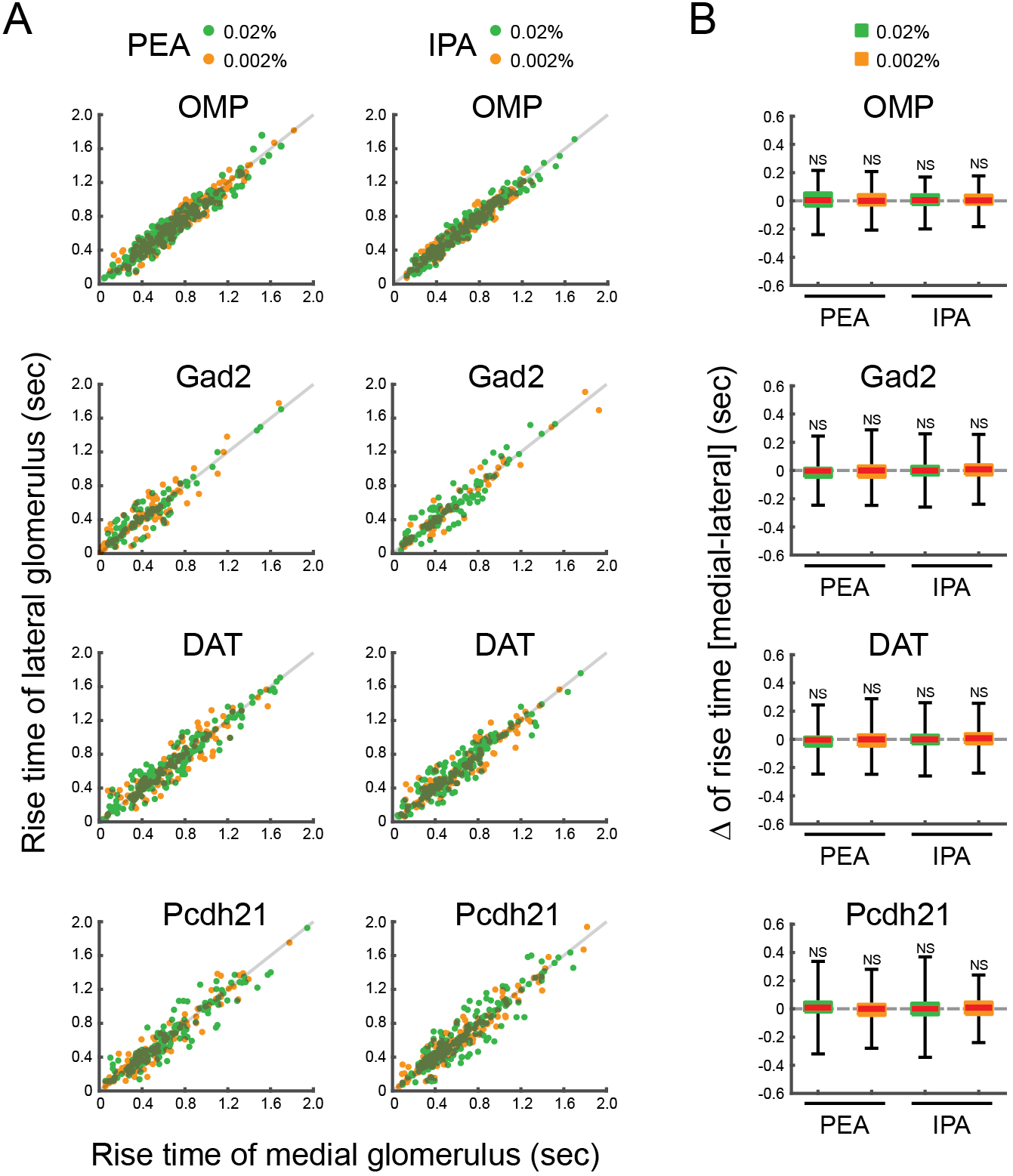
Rise times of medial and lateral glomeruli. (A) Scatter plots displaying the distributions of rise times of the medial and lateral glomeruli, which responded to PEA and IPA stimuli. x and y axes indicate rise times of medial and lateral glomerular responses, respectively. Individual green and orange dots indicate single trial data of 0.02% and 0.002% PEA or IPA. Gray lines indicate the equal rise time points of medial and lateral glomerular responses. (B) Box plots displaying the distributions of the differences of rise times between medial and lateral glomerular responses to 0.02% and 0.002% PEA or IPA. Red horizontal lines in each box indicate the medians. Quartiles are shown as whiskers. NS, not significant (two-tailed paired Student’s t test).

**Figure 6:**
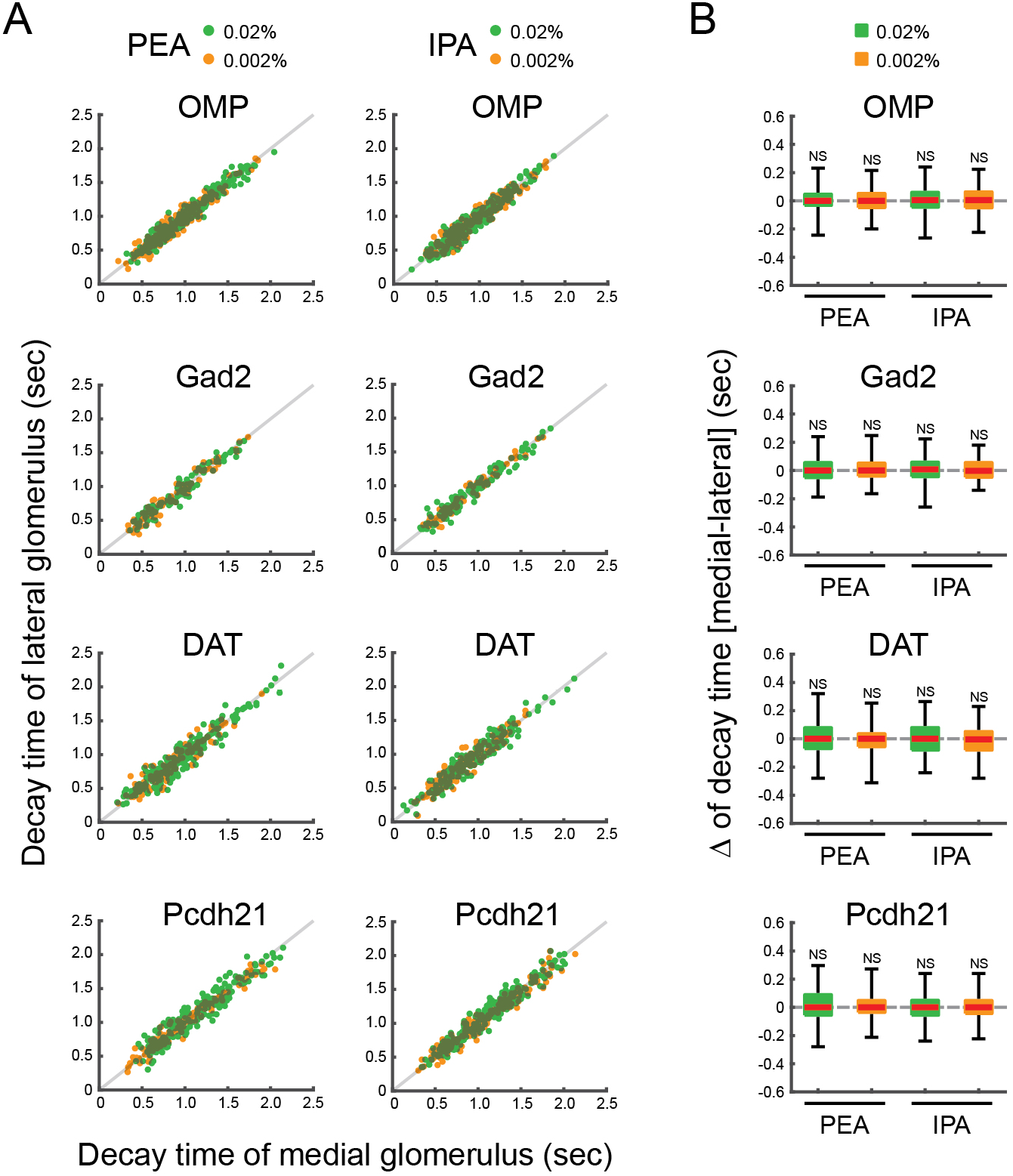
Decay times of medial and lateral glomeruli. (A) Scatter plots displaying the distributions of decay times of the medial and lateral glomeruli, which responded to PEA and IPA stimuli. x and y axes indicate decay times of medial and lateral glomerular responses, respectively. Individual green and orange dots indicate single trial data of 0.02% and 0.002% PEA or IPA. Gray lines indicate the equal decay time points of medial and lateral glomerular responses. (B) Box plots displaying the distributions of the differences of decay times between medial and lateral glomerular responses to 0.02% and 0.002% PEA or IPA. Red horizontal lines in each box indicate the medians. Quartiles are shown as whiskers. NS, not significant (two-tailed paired Student’s t test).

**Figure 7:**
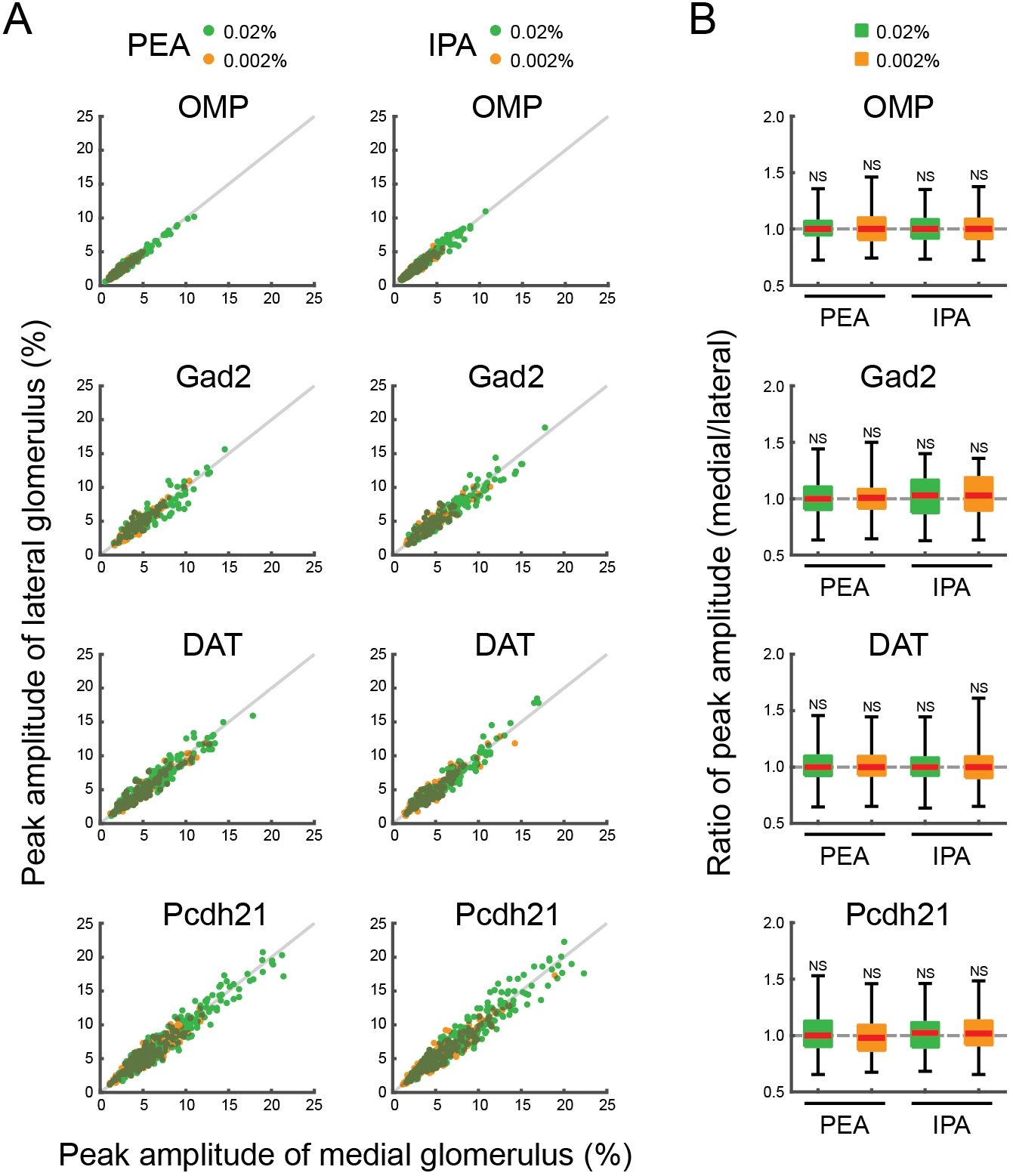
Peak amplitude of medial and lateral glomerular responses. (A) Scatter plots displaying the distributions of peak amplitudes of medial and lateral glomerular responses to PEA and IPA stimuli. x and y axes indicate peak amplitudes of medial and lateral glomerular responses, respectively. Individual green and orange dots indicate single trial data of 0.02% and 0.002% PEA or IPA. Gray lines indicate the equal peak amplitudes of medial and lateral glomerular responses. (B) Box plots displaying the distributions of the differences of peak amplitudes of medial and lateral glomerular responses to 0.02% and 0.002% PEA or IPA. Red horizontal lines in each box indicate the medians. Quartiles are shown as whiskers. NS, not significant (two-tailed paired Student’s t test).

**Table 1.**
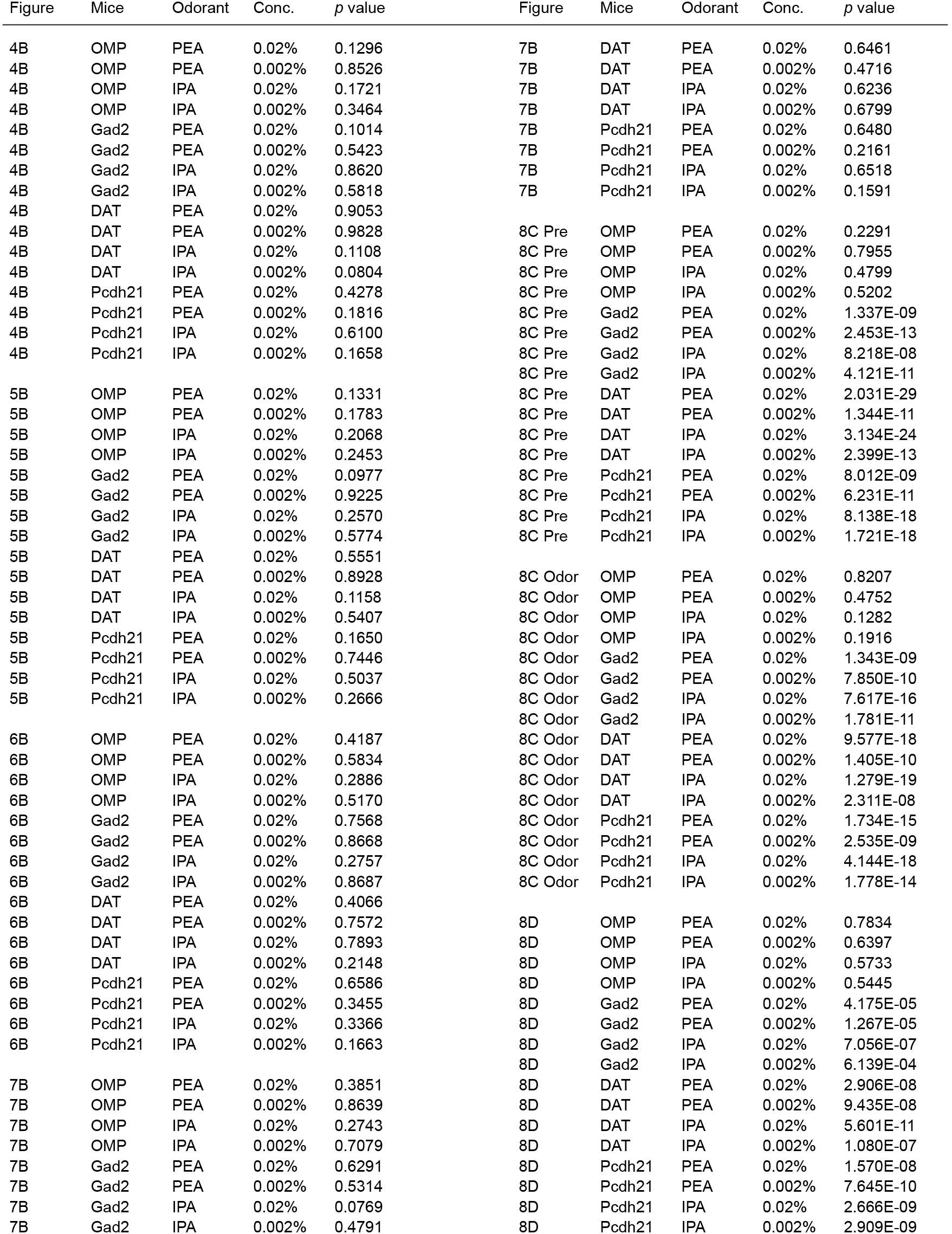

| Figure | Mice | Odorant | Conc. | p value | Figure | Mice | Odorant | Conc. | p value |
| --- | --- | --- | --- | --- | --- | --- | --- | --- | --- |
| 4B | OMP | PEA | 0.02% | 0.1296 | 7B | DAT | PEA | 0.02% | 0.6461 |
| 4B | OMP | PEA | 0.002% | 0.8526 | 7B | DAT | PEA | 0.002% | 0.4716 |
| 4B | OMP | IPA | 0.02% | 0.1721 | 7B | DAT | IPA | 0.02% | 0.6236 |
| 4B | OMP | IPA | 0.002% | 0.3464 | 7B | DAT | IPA | 0.002% | 0.6799 |
| 4B | Gad2 | PEA | 0.02% | 0.1014 | 7B | Pcdh21 | PEA | 0.02% | 0.6480 |
| 4B | Gad2 | PEA | 0.002% | 0.5423 | 7B | Pcdh21 | PEA | 0.002% | 0.2161 |
| 4B | Gad2 | IPA | 0.02% | 0.8620 | 7B | Pcdh21 | IPA | 0.02% | 0.6518 |
| 4B | Gad2 | IPA | 0.002% | 0.5818 | 7B | Pcdh21 | IPA | 0.002% | 0.1591 |
| 4B | DAT | PEA | 0.02% | 0.9053 |  |  |  |  |  |
| 4B | DAT | PEA | 0.002% | 0.9828 | 8C Pre | OMP | PEA | 0.02% | 0.2291 |
| 4B | DAT | IPA | 0.02% | 0.1108 | 8C Pre | OMP | PEA | 0.002% | 0.7955 |
| 4B | DAT | IPA | 0.002% | 0.0804 | 8C Pre | OMP | IPA | 0.02% | 0.4799 |
| 4B | Pcdh21 | PEA | 0.02% | 0.4278 | 8C Pre | OMP | IPA | 0.002% | 0.5202 |
| 4B | Pcdh21 | PEA | 0.002% | 0.1816 | 8C Pre | Gad2 | PEA | 0.02% | 1.337E-09 |
| 4B | Pcdh21 | IPA | 0.02% | 0.6100 | 8C Pre | Gad2 | PEA | 0.002% | 2.453E-13 |
| 4B | Pcdh21 | IPA | 0.002% | 0.1658 | 8C Pre | Gad2 | IPA | 0.02% | 8.218E-08 |
|  |  |  |  |  | 8C Pre | Gad2 | IPA | 0.002% | 4.121E-11 |
| 5B | OMP | PEA | 0.02% | 0.1331 | 8C Pre | DAT | PEA | 0.02% | 2.031E-29 |
| 5B | OMP | PEA | 0.002% | 0.1783 | 8C Pre | DAT | PEA | 0.002% | 1.344E-11 |
| 5B | OMP | IPA | 0.02% | 0.2068 | 8C Pre | DAT | IPA | 0.02% | 3.134E-24 |
| 5B | OMP | IPA | 0.002% | 0.2453 | 8C Pre | DAT | IPA | 0.002% | 2.399E-13 |
| 5B | Gad2 | PEA | 0.02% | 0.0977 | 8C Pre | Pcdh21 | PEA | 0.02% | 8.012E-09 |
| 5B | Gad2 | PEA | 0.002% | 0.9225 | 8C Pre | Pcdh21 | PEA | 0.002% | 6.231E-11 |
| 5B | Gad2 | IPA | 0.02% | 0.2570 | 8C Pre | Pcdh21 | IPA | 0.02% | 8.138E-18 |
| 5B | Gad2 | IPA | 0.002% | 0.5774 | 8C Pre | Pcdh21 | IPA | 0.002% | 1.721E-18 |
| 5B | DAT | PEA | 0.02% | 0.5551 |  |  |  |  |  |
| 5B | DAT | PEA | 0.002% | 0.8928 | 8C Odor | OMP | PEA | 0.02% | 0.8207 |
| 5B | DAT | IPA | 0.02% | 0.1158 | 8C Odor | OMP | PEA | 0.002% | 0.4752 |
| 5B | DAT | IPA | 0.002% | 0.5407 | 8C Odor | OMP | IPA | 0.02% | 0.1282 |
| 5B | Pcdh21 | PEA | 0.02% | 0.1650 | 8C Odor | OMP | IPA | 0.002% | 0.1916 |
| 5B | Pcdh21 | PEA | 0.002% | 0.7446 | 8C Odor | Gad2 | PEA | 0.02% | 1.343E-09 |
| 5B | Pcdh21 | IPA | 0.02% | 0.5037 | 8C Odor | Gad2 | PEA | 0.002% | 7.850E-10 |
| 5B | Pcdh21 | IPA | 0.002% | 0.2666 | 8C Odor | Gad2 | IPA | 0.02% | 7.617E-16 |
|  |  |  |  |  | 8C Odor | Gad2 | IPA | 0.002% | 1.781E-11 |
| 6B | OMP | PEA | 0.02% | 0.4187 | 8C Odor | DAT | PEA | 0.02% | 9.577E-18 |
| 6B | OMP | PEA | 0.002% | 0.5834 | 8C Odor | DAT | PEA | 0.002% | 1.405E-10 |
| 6B | OMP | IPA | 0.02% | 0.2886 | 8C Odor | DAT | IPA | 0.02% | 1.279E-19 |
| 6B | OMP | IPA | 0.002% | 0.5170 | 8C Odor | DAT | IPA | 0.002% | 2.311E-08 |
| 6B | Gad2 | PEA | 0.02% | 0.7568 | 8C Odor | Pcdh21 | PEA | 0.02% | 1.734E-15 |
| 6B | Gad2 | PEA | 0.002% | 0.8668 | 8C Odor | Pcdh21 | PEA | 0.002% | 2.535E-09 |
| 6B | Gad2 | IPA | 0.02% | 0.2757 | 8C Odor | Pcdh21 | IPA | 0.02% | 4.144E-18 |
| 6B | Gad2 | IPA | 0.002% | 0.8687 | 8C Odor | Pcdh21 | IPA | 0.002% | 1.778E-14 |
| 6B | DAT | PEA | 0.02% | 0.4066 |  |  |  |  |  |
| 6B | DAT | PEA | 0.002% | 0.7572 | 8D | OMP | PEA | 0.02% | 0.7834 |
| 6B | DAT | IPA | 0.02% | 0.7893 | 8D | OMP | PEA | 0.002% | 0.6397 |
| 6B | DAT | IPA | 0.002% | 0.2148 | 8D | OMP | IPA | 0.02% | 0.5733 |
| 6B | Pcdh21 | PEA | 0.02% | 0.6586 | 8D | OMP | IPA | 0.002% | 0.5445 |
| 6B | Pcdh21 | PEA | 0.002% | 0.3455 | 8D | Gad2 | PEA | 0.02% | 4.175E-05 |
| 6B | Pcdh21 | IPA | 0.02% | 0.3366 | 8D | Gad2 | PEA | 0.002% | 1.267E-05 |
| 6B | Pcdh21 | IPA | 0.002% | 0.1663 | 8D | Gad2 | IPA | 0.02% | 7.056E-07 |
|  |  |  |  |  | 8D | Gad2 | IPA | 0.002% | 6.139E-04 |
| 7B | OMP | PEA | 0.02% | 0.3851 | 8D | DAT | PEA | 0.02% | 2.906E-08 |
| 7B | OMP | PEA | 0.002% | 0.8639 | 8D | DAT | PEA | 0.002% | 9.435E-08 |
| 7B | OMP | IPA | 0.02% | 0.2743 | 8D | DAT | IPA | 0.02% | 5.601E-11 |
| 7B | OMP | IPA | 0.002% | 0.7079 | 8D | DAT | IPA | 0.002% | 1.080E-07 |
| 7B | Gad2 | PEA | 0.02% | 0.6291 | 8D | Pcdh21 | PEA | 0.02% | 1.570E-08 |
| 7B | Gad2 | PEA | 0.002% | 0.5314 | 8D | Pcdh21 | PEA | 0.002% | 7.645E-10 |
| 7B | Gad2 | IPA | 0.02% | 0.0769 | 8D | Pcdh21 | IPA | 0.02% | 2.666E-09 |
| 7B | Gad2 | IPA | 0.002% | 0.4791 | 8D | Pcdh21 | IPA | 0.002% | 2.909E-09 |

### Respiration-locked calcium fluctuations in medial maps are larger than in lateral maps

Oscillatory calcium responses associated with respiration were observed in the optical recordings (Fig. 3B). These respiratory-linked fluctuations in calcium signals in medial glomeruli appeared to be larger in the postsynaptic neurons (i.e., GABAergic, dopaminergic, and mitral/tufted cells in the Gad2-Cre, DAT-Cre, and Pcdh21-Cre lines, respectively) than in the OSNs (i.e., cells in the OMP-Cre line). To examine this quantitatively, we applied a power spectral analysis to the data. Power spectra before and during odor stimulation displayed peak frequencies of 2–4 Hz (arrows in Fig. 8A), which matches the respiration rhythm under our experimental conditions. These respiration-locked calcium oscillations were detected in all the cell types (Fig. 8B). The peak power spectra at 2–4 Hz were indeed larger in medial glomeruli than in lateral glomeruli from Gad2-, DAT-, and Pcdh21-Cre mice both before and after odor stimulation but not in OMP-Cre mice. The statistical analysis is summarized in Fig. 8C (two-tailed paired *t* tests, see Table 1; the datasets are the same as in Fig. 4–7). In addition, the medial/lateral power ratios for spectra from postsynaptic neurons were larger during odor stimulation than before stimulation (Fig. 8D), suggesting a strong influence of the odor stimulus on the size of the fluctuation. These differences were observed consistently with 0.02% and 0.002% PEA and IPA (two-tailed paired *t* tests, see Table 1). These results suggest that respiration-locked calcium fluctuations in the medial maps are enhanced postsynaptically in the OB by odorant stimulation.

**Figure 8:**
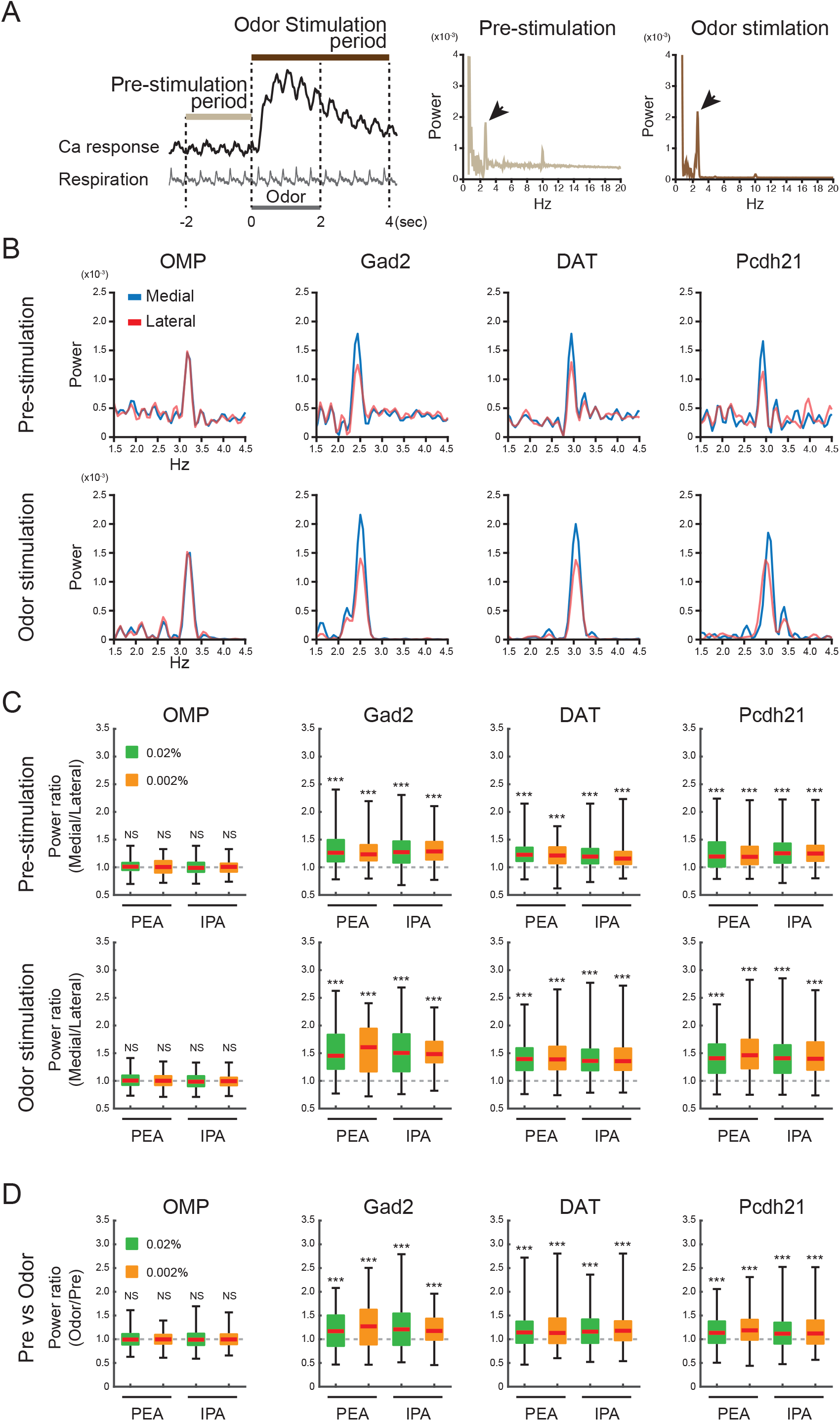
Respiration-locked calcium fluctuations of medial and lateral glomeruli. (A) Schematic illustration of data collection time points and power spectral analysis. The prestimulation and odor stimulation periods are defined in left panel. Middle and right panels are representative examples of power spectral analyses with large frequency ranges in single trials. Respiration-locked fluctuations of calcium signals were prominently observed at 2–4 Hz (arrows). (B) Representative results of power spectral analyses of calcium signals during prestimulation (upper panels) and odor stimulation (lower panels) periods in neurons from GCaMP-expressing OMP-, Gad2-, DAT-, and Pcdh21-Cre mice. These calcium response traces are shown as 0.02% IPA response signals (as in Fig. 3B). Blue and red traces indicate medial and lateral glomeruli, respectively. (C) Comparisons of power spectra between the homologous glomeruli during the prestimulation (upper panels) and odor stimulation (lower panels) period in neurons from OMP-, Gad2-, DAT-, and Pcdh21-Cre mice. The ratios of peak powers were calculated by dividing the medial glomerular power by the lateral glomerular power in each trial. (D) Comparisons of peak power between the prestimulation and odor stimulation periods. The power ratios were calculated by dividing the odor stimulation period power (lower panels in C) by the prestimulation-period power (show in C, upper panels). Red horizontal lines in each box show medians. Quartiles are shown as whiskers. NS, not significant; ***; p < 0.001 two-tailed paired Student’s t test).

## Discussion

### Pre/postsynaptic calcium events in multiple cell types

In this work, GCaMP was expressed in postsynaptic GABAergic, dopaminergic, and mitral/tufted cells and in presynaptic axon terminals of OSNs of mice from various transgenic Cre-driver lines. Calcium signaling in OSNs reflects activation resulting in transmitter release, whereas calcium signaling in the other cell types may reflect activation of calcium-permeable glutamate receptors and the opening of calcium channels in response to excitatory postsynaptic potentials or spikes initiated in the dendrites or soma (Burnashev et al., 1992; Chen et al., 1997; Halabisky et al., 2000; Helmchen et al., 1999; Nagayama et al., 2007; Svoboda et al., 1999). Therefore, the calcium influxes in these cells are controlled by different biophysical mechanisms and represent different aspects of biological events. These differences would not impact our imaging results, as the comparisons were between medial and lateral glomeruli comprising the same cell types. Notably, we did not observe cell-type-specific differences between the medial and lateral maps at the glomerular level, which is consistent with recent data at the single-cell level showing that odorant responses of juxtaglomerular cells are associated with the same glomerulus putatively (Homma et al., 2019). Nevertheless, we cannot exclude the possibility of differential response timing by different cell types, because the time resolution for calcium imaging was limited and did not reflect that activity of all neurons within a glomerulus. Future studies may begin to address this by imaging spike activity at a single-cell resolution.

### Response timing of the homologous glomeruli in medial and lateral maps

A pioneer electrophysiological study using a unique transgenic mouse line in which all OSNs express the same odorant receptor suggested that that the latencies of mitral cell responses to odorant stimulation are shorter in the medial map than in the lateral map, especially at a low odorant concentration (Zhou and Belluscio, 2012). Because the OE is a complicated structure, airflow speed likely varies throughout the nasal cavity, and odorants may reach different areas at different times (Kimbell et al., 1997; Schoenfeld and Cleland, 2006; Zhao et al., 2006). This would produce a time lag for odorant responses in medial and lateral neurons. However, we did not observe different onset latencies between homologous medial and lateral glomeruli, even at low odorant concentrations. This may be because our recordings were via optical imaging rather than electrophysiology and from glomerular rather than MCLs. Another possible reason is the difference in OB regions recorded, which reflect inputs from different OE regions. Our study was restricted to a small region of the posteromedial dorsal OB, which receives inputs from the dorsal OE (Miyamichi et al., 2005). The dorsal OE faces the large nasal cavity and has a relatively simple wall structure compared with that of ventral zones. Moreover, OSNs projecting to medial and lateral maps are close together in the dorsal OE, for which any latency would not be detectable under our experimental conditions. It is possible that differential latencies from neurons in the ventral OB may be larger or more easily detected (Kimbell et al., 1997; Schoenfeld and Cleland, 2006; Zhao et al., 2006). In other words, the time lag may gradually become larger along the dorsal-ventral axis in the OB.

### Neuronal/circuitry mechanism of respiration-locked calcium fluctuations

The fluctuations of calcium responses, which corresponded to the rhythm of respiration, were larger in medial glomeruli than in lateral glomeruli. This may reflect differential airflow volumes or rates along the medial and lateral sides of the nostril. As this was only observed in postsynaptic calcium responses, the modulation is likely not within the OE but in the OB. The mechanism for this modulation may involve the physiological properties of these and/or associated neurons in medial and lateral glomeruli, such as differences in the expression of various calcium and/or other essential channels. As we did not observe differences in the amplitudes of the calcium responses, more than one channel type may be involved. Differential expression of other essential molecules would also change neuronal and/or network excitability. Neurons in medial glomeruli may also increase and decrease their intracellular calcium levels more synchronously during inhalation and exhalation, respectively. Such synchrony would be expected to affect the overall activity of neurons within a glomerulus; this could more directly be addressed by studies recording neuronal spikes in the context of a circuit. Thus, further investigations are needed to determine the mechanism by which calcium fluctuations are larger in medial glomeruli than in lateral glomeruli in response to odorant stimulation.

### Inhibitory connections between the two maps

Tufted cells in the lateral glomeruli project to cells in the internal plexiform and superficial GCLs underlying the medial glomeruli receiving inputs from the same odorant receptors, and vice versa. These projections activate granule cells and thus inhibit surrounding mitral/tufted cells (Belluscio and Katz, 2001; Lodovichi et al., 2003; Zhou and Belluscio, 2008), resulting in mutual inhibition between the medial and lateral maps. Our data suggest this inhibition is not simple (i.e., one glomerulus is inhibited when the other is activated), as medial and lateral glomeruli are activated simultaneously during odor stimulation. The similar temporal patterns and the absence of counterphase-locked activity between the two glomeruli also suggest that the activity of one glomerulus does not inhibit the other homologous glomerulus in a given time phase, such as during inhalation or exhalation. The functional role of these mutual inhibitory connections and how they contribute to odor processing remain unknown. It is possible that these connections regulate the temporal activity pattern and/or synchrony of neurons or glomeruli in both maps.

### Odor information processing streams from the medial and lateral maps

One of the unresolved issues is where the medial and lateral maps project and how higher brain centers handle these two outputs. Current knowledge of the connections between the OB and the olfactory cortex (Ghosh et al., 2011; Igarashi et al., 2012; Miyamichi et al., 2011; Nagayama et al., 2010; Sosulski et al., 2011) suggest that there may not be dramatic differences regarding where the maps project. However, it is still not known whether the outputs from the two maps are evenly transmitted to all olfactory cortical areas, and some regions may preferentially receive input from one or the other. More detailed knowledge of the cortical projections may help reveal the significance of the multiple maps and how the information from the two information streams is compiled in higher brain centers.

### Rhythm of respiration

In vertebrate land animals, airflow through the nasal cavity during respiration alternates between orthonasal and retronasal directions. This induces a synchronized rhythm in the olfactory system, which contributes to the odor information process (Cury and Uchida, 2010; Spors and Grinvald, 2002; Uchida et al., 2014; Wilson and Mainen, 2006). Moreover, the orthonasal and retronasal airflows switch the perception from smells originating from the surrounding environment to taste in the mouth, respectively (Gautam and Verhagen, 2012; Shepherd, 2012). In addition to olfactory areas, respiration-locked oscillations have been observed in hippocampus as well as barrel and prefrontal cortices in the rodent (Biskamp et al., 2017; Lockmann et al., 2016; Nguyen Chi et al., 2016; Phillips et al., 2012; Shusterman et al., 2011; Yanovsky et al., 2014). In the barrel cortex, phase-locked oscillation patterns coordinate the interaction between olfaction and tactile sensations (Ito et al., 2014), whereas freezing behavior is modulated by rhythmic activity in prelimbic prefrontal cortex driven by inputs from the anterior olfactory nucleus (Moberly et al., 2018), which has topographical connections to the OB (Schoenfeld et al., 1985; Yan et al., 2008). The anterior olfactory nucleus sub-region which dominantly primarily receive inputs from the medial map may relay this information to higher brain centers controlling respiration-linked neuronal activity associated with mouse behavior. Thus, the anterior olfactory nucleus may be responsible for processing information from multiple glomerular maps and represents an area that warrants further study.

## Materials and Methods

All procedures were performed on mice of either sex in accordance with National Institutes of Health guidelines and approved by the Animal Welfare Committee at the University of Texas Health Science Center at Houston.

### Animals

Cre-inducible GCaMP3-expressing mice (Ai38; #014538, Jackson Laboratory, Bar Harbor, ME) (Zariwala et al., 2012) were used for the expression of the calcium indicator in target neurons. This mouse line was crossed with the following Cre-driver mouse lines: OMP-Cre (for OSNs, #006668; Jackson Laboratory) (Li et al., 2004), Gad2-Cre (for GABAergic neurons, #010802; Jackson Laboratory) (Taniguchi et al., 2011), DAT-Cre (for dopaminergic neurons, #006660; Jackson Laboratory) (Bäckman et al., 2006), and Pcdh21-Cre (for mitral/tufted cells, #02189; RIKEN BioResource Research Center, Tsukuba, Japan) (Nagai et al., 2005).

### Histology

Mice were deeply anesthetized and fixed by transcardial perfusion with 4% paraformaldehyde (PFA) in 0.1 M phosphate buffer (PB; pH 7.4). Then, whole brains were dissected out and postfixed in 4% PFA/0.1 M PB overnight. The samples were cryoprotected in 30% sucrose (wt/vol) in phosphate-buffered saline (PBS; pH 7.4) and embedded in optimal cutting temperature compound (Fisher HealthCare, Waltham, MA). Olfactory tissue sections 30-μm thick were cut on a cryostat, washed with PBS, and mounted with Fluoroshield mounting medium (F6057; Sigma-Aldrich, St. Louis, MO). Images were captured on an Olympus FluoView FV1000 laser scanning confocal microscope using a 20×/0.75 NA lens objective (UPLSAPO 20X; Olympus, Tokyo, Japan).

### Odorant stimulation

IPA (#126810; Sigma-Aldrich) and PEA (#128945; Sigma-Aldrich) were diluted in mineral oil (M3516; Sigma-Aldrich) to 0.1% and 0.01% in glass vials. The odorants were vaporized using nitrogen, mixed with filtered air to final concentrations of 0.02% and 0.002% with 0.5 liter/min air flow rate, and then delivered to mouse nostrils using a custom-made olfactometer (Kikuta et al., 2013). The odorants were presented for 2 s with an interstimulus interval of >60 s to avoid sensory adaptation.

### *In vivo* optical imaging

GCaMP-expressing mice were anesthetized with urethane (1.2 g/kg of body weight, intraperitoneal). The depth of anesthesia was monitored by toe pinches throughout the experiment. Body temperature was kept between 36.0°C and 37.0°C with a heating pad. The skull over the OBs was carefully thinned with a dental drill and covered with 1.2% agarose dissolved in saline and with a 4–6-mm^2^ coverslip (thickness, #1). Odor-evoked GCaMP3 signals were recorded through a 5× lens objective (Fluar 5×/0.25; Zeiss, Oberkochen, Germany) on a microscope (SliceScope; Scientifica, Uckfield, United Kingdom) equipped with a high-speed charge-coupled-device camera (NeuroCCD-SM256; RedShirtImaging, Decatur, GA) at 125 Hz (128 × 128 pixels) for 12 s, which included a 4-s prestimulus period and a 2-s odor presentation period. Excitation light was provided using a 470-nm light-emitting diode module (M470L2; Thorlabs, Newton, NJ). A standard green fluorescent protein filter set (BrightLine GFP-4050A-OMF-ZERO; Semrock, Rochester, NY) was used to detect the GCaMP3 signal. Chest movement of the animals was monitored to measure the respiratory rhythm during the optical imaging period.

### DiI labeling of OSN axons

After calcium imaging, the skull over the dorsal OBs was removed. Then, small DiI crystals were attached to the tip of glass capillaries (tip diameter, ~5 μm) and embedded into the area of IPA-responsive glomeruli. The positions of blood vessels relative to identical pairs of glomeruli were used as landmarks for DiI implantation. The mice remained anesthetized for 9–10 h after DiI implantation and then were fixed by transcardial perfusion with 4% PFA/0.1 M PB. Whole brains were removed and incubated in PBS at room temperature until observation. Low-magnification images were captured by a charge-coupled-device camera (NeuroCCD-SM256 or SensiCam; PCO, Kelheim,Germany) using a 5× lens objective (Fluar 5×/0.25, Zeiss) with a light-emitting diode module (MCWHL2-C1; Thorlabs) and a standard Cy3 cube (BrightLine Cy3-4040C-OMF-ZERO; Semrock). For high-magnification views of OSN axons, images were acquired with a two-photon microscope (Prairie Ultima; Bruker, Billerica, MA), using a 20× water immersion lens objective (UMPLFLH 20XW; Olympus), with a 5-μm inter-z-slice interval and 512 × 512 pixel resolution. DiI was excited at 920 nm (Ti:sapphire laser, MaiTai HP DS; Spectra-Physics, Santa Clara, CA), and DiI fluorescence was detected with an emission filter cube (575-nm dichroic mirror and 607/45-nm barrier filters).

### Data analysis for wide-field calcium imaging data

Odor-evoked response maps were generated using Fiji/ImageJ (Schindelin et al., 2012) with custom-written scripts. Spatially filtered (3 × 3 mean filter) prestimulation-period images (4 s) were averaged and used as a baseline (F_0_). Images averaged 3 s after the onset of the 1-s odor stimulation were used as the response image (F). Then, F was subtracted from F_0_ to obtain the difference (ΔF). ΔF values were divided by F_0_ to obtain the ratio image (ΔF/F_0_). Spatial filters (3 × 3 mean filter) were also applied to the ratio images. All negative values were set to zero in the images. Regions of interest corresponding to glomeruli were manually set (4–8 pixels centered on each glomeruli).

The time courses of calcium signals (see Fig. 3B) were calculated using MATLAB (MathWorks, Natick, MA) with custom-written scripts. ΔF/F_0_ values were calculated using the same procedure described above but with a temporal filter (3 frames average, 24 ms) rather than a spatial filter applied to the ΔF/F_0_ values.

Onset latency, rise time, decay time, and peak amplitude were computed with custom MATLAB scripts. First, the baseline values (ΔF/F_0_) were determined as the mean values over the baseline period, which were defined as the 80-ms time window immediately before stimulation onset. Then, the noise level in each trial was defined as the minimum standard deviation among eight 0.5-s blocks in the 4-s prestimulation period. The onset latency was determined as the first time point at which all data points in the subsequent 80 ms exceeded the threshold (2.5 times the noise level). The onset latency was measured as the time elapsed from the first inhalation after the onset of odor presentation. The rise time was defined as the duration the calcium signals increased from 20% to 80% of peak amplitude. Time points for when the calcium signals reached 20% and 80% of the peak amplitude were set as the earliest time point after which half of the data points in the subsequent 80 ms exceeded these criteria. The decay time was defined as the duration the calcium signal decreased from 100% to 50% of the peak amplitude. Time points for when the calcium signals reached 100% and 50% of peak amplitude were set as the earliest time point after which half of data points in the subsequent 80 ms dropped below these criteria. The peak amplitude was measured as the maximum value using the 80-ms time window moving average, which reflects average of 9 sequential data points, after stimulation onset.

To analyze fluctuations in the calcium fluorescence, a power spectral analysis based on Fourier transform was used. First, ΔF/F_0_ values were preprocessed with a 40-ms box filter and divided into two periods, corresponding to prestimulation (2 s before onset of odor stimulation) and odor stimulation (4 s after onset of odor stimulation). Then, a power spectrum was computed by using Fast Fourier transform with 2,048 points (using a built-in function of MATLAB [fft.m]) in each period. Because we used a 125-Hz sampling frequency, the frequency resolution is 0.061 Hz. The peak power in each period was determined as the maximum value between 2 and 4 Hz.

### Statistics

Statistical analyses were performed using Microsoft Excel 2013. All statistical significance was determined by a two-tailed paired Student’s *t* test; *p* values of <0.05 were considered statistically significant. Data are presented as scatter and box-whisker plots of pooled data sets for a given odorant and concentration from OMP- (279 trials in 0.02% PEA, 194 trials in 0.002% PEA, 297 trials in 0.02% IPA, and 185 trials in 0.002% IPA; *n* = 10), Gad2- (106 trials in 0.02% PEA, 84 trials in 0.002% PEA, 119 trials in 0.02% IPA, and 62 trials in 0.002% IPA; *n* = 6), DAT- (216 trials in 0.02% PEA, 118 trials in 0.002% PEA, 184 trials in 0.02% IPA, and 123 trials in 0.002% IPA; *n* = 6), and Pcdh21-Cre (199 trials in 0.02% PEA, 130 trials in 0.002% PEA, 241 trials in 0.02% IPA, and 176 trials in 0.002% IPA; *n* = 6) mice. In scatter plots, individual dots show data points from single trials. In box-whisker plots, horizontal red lines and boxes indicate the medians and quartiles, respectively. The whiskers go from each quartile to the minimum or maximum. All *p* values calculated in this study are listed in Table 1.

## Acknowledgements

We thank Dr. Dewan for discussions of odorant response properties and the location of TAAR glomeruli.

